# Abstract Encoding of Sounds in the Frontopolar Cortex

**DOI:** 10.64898/2026.02.23.707465

**Authors:** Marlen Alva, José Vergara, Tonatiuh Figueroa, Luis Lemus

## Abstract

The frontopolar cortex has been linked to higher-order cognition, including analogical reasoning, cost-benefit analysis, and semantic associations, primarily based on neuroimaging studies in humans rather than from direct single-neuron recordings. Notably, sensory input to this region originates from auditory cortical areas. To investigate the role of the frontopolar cortex in auditory perception, we recorded single neurons in nonhuman primates trained to discriminate between various sounds, including monkey calls and human words. We found that individual neurons form abstract, nonlinear representations of learned and novel sounds, collectively encoding all sound categories and generating decision-making signals. Our findings suggest that the frontopolar cortex integrates auditory information into behaviorally relevant signals, thereby providing insights regarding its role in cognition.

## Introduction

The frontopolar cortex (FPC), also known as Brodmann’s area 10 (BA10), is a cortical region in the primate brain (Preuss & Goldman-Rakic, 1991) that shows substantial expansion in humans (Ongur & Price, 2000; Semendeferi et al., 2001; Tsujimoto et al., 2011; Mansouri et al., 2017). Neuroimaging studies indicate that the FPC is critical for high-level cognitive functions (Ramnani & Owen, 2004), including decision-making based on cost-benefit analysis, the management of ongoing and prospective goals, and the switching between exploration and exploitation (Tsujimoto et al., 2011; Mansouri et al., 2014; 2017; Koechlin et al., 1999; Pollmann, 2016; Sakai, 2008; Dreher et al., 2008; Gilbert et al., 2005; 2006; Zajkowski et al., 2017; Boorman et al., 2009). It has also been associated with sequence assembly (Desrochers et al., 2015, 2019) and episodic memory (Kim, 2013). Notably, FPC ablations in monkeys have shown improved performance on conflicting decisions (Mansouri et al., 2015), but impair the learning of novel stimuli (Boschin et al., 2015; Miyamoto et al., 2018).

The FPC has a significant role in abstract thinking (Amodio & Frith, 2004; Koechlin & Hyafil, 2007; Badre & D’Esposito, 2009; Bengston et al., 2009) and analogical associations in language (Klein et al., 1995; Braver & Bongolatti, 2002; Green et al., 2006, 2010; Bunge et al., 2005; 2009), which probably arise from the FPC’s proposed specialization in auditory processing (Medalla & Barbas, 2014). In fact, FPC’s densest connections are with auditory regions in the superior temporal gyrus (STG; Burman et al., 2011; Petrides & Pandya, 1988; 2007; Barbas & Mesulam, 1985; Barbas et al., 1999; Hackett et al., 1999). Furthermore, it lacks direct visual inputs from the occipital, parietal, or inferotemporal cortices.

Despite FPC connectivity, nonhuman primates performing on visual paradigms are among the few studies that have recorded single neurons directly and have shown that these neurons respond to uncertainty and feedback (Nougaret et al., 2024; Ferrucci et al., 2022; Tsujimoto et al., 2010). Therefore, to address how FPC contributes to auditory perception, we performed single-unit recordings in two rhesus monkeys trained to discriminate between learned sounds such as conspecific vocalizations and words (Melchor et al., 2021; Morán et al., 2021). We found that, while single neurons create abstract representations of a limited set of learned and novel sounds, the neural population encodes the complete set and generates decision-making signals.

## Results

### Neuronal coding of auditory categories

To determine the role of the frontopolar cortex (Fig. 1a) in auditory processing, we performed single-unit recordings in two rhesus macaques discriminating animal vocalizations and words learned as targets or distractors (Fig. 1b). In each trial, a sequence of sounds was delivered, each followed by a short delay and a visual go-cue window (Fig. 1c). Trials included one or more sounds randomly selected from the groups of targets and distractors, with a target always presented last (Fig. 1d). The animals received a liquid reward for hits, releasing the lever only during the go-cue window following a target (Fig. 1e).

**Fig. 1.**
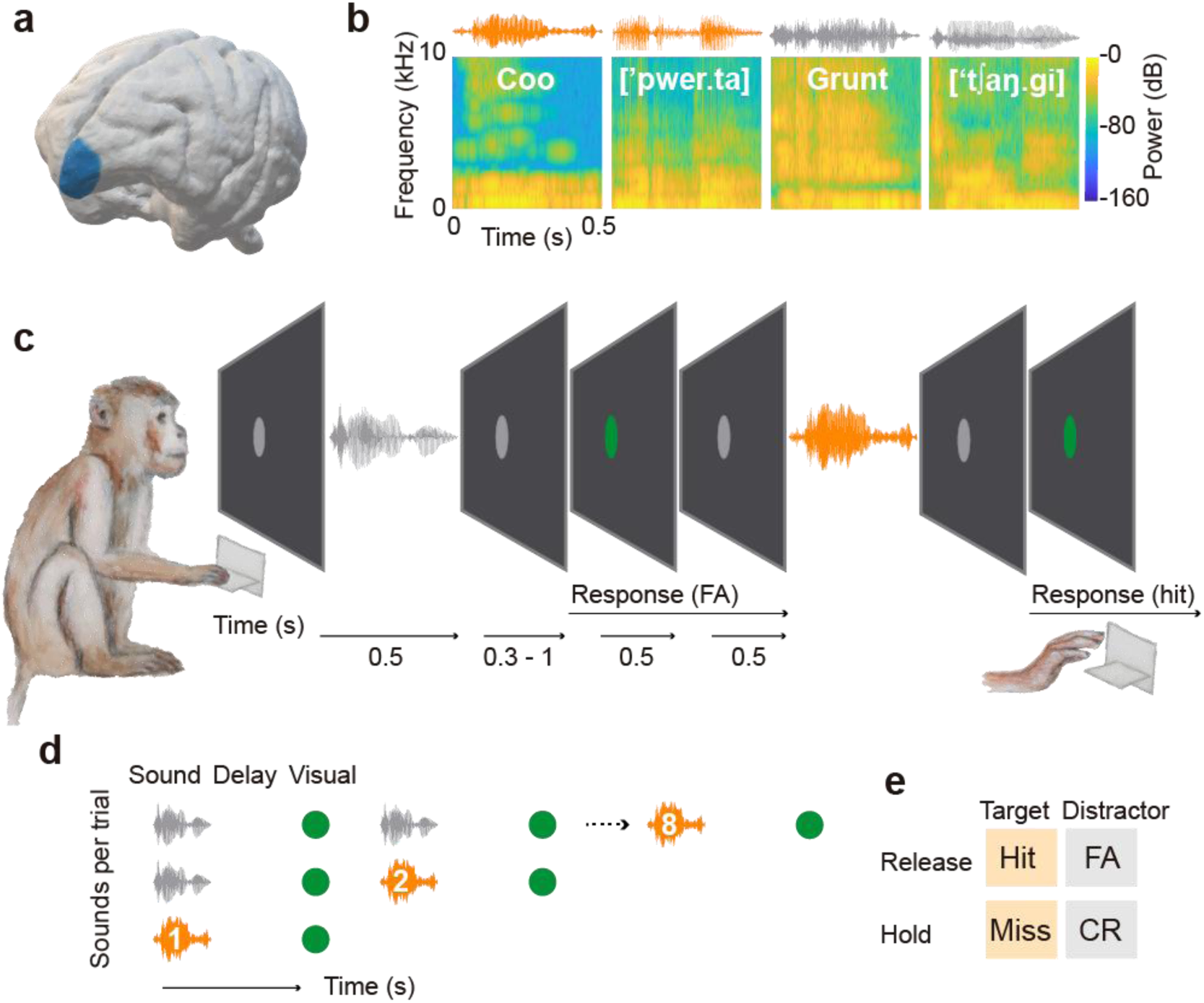
The frontopolar cortex and behavioral paradigm. **a**, The left FPC is depicted on a structural MRI scan of monkey V. **b**, Sonograms and spectrograms illustrate representative target (orange) and distractor (gray) sounds utilized in the auditory discrimination task. Spanish words are transcribed using the International Phonetic Alphabet. **c**, The trial sequence began when monkeys pressed and held a lever in response to a gray circle displayed on the screen, initiating a sequence of 1–3 sounds (0.5 s each) in monkey V or 1–7 sounds (0.5 s each) in monkey X. Each sound was followed by a delay (0.3-0.6 s in monkey V, 1 s in monkey X), then a 0.5 s go-cue period during which the gray circle turned green to signal the onset of a 1 s response window. Monkeys were required to maintain lever hold during distractor periods to achieve a correct rejection (CR), and to release the lever only within a target response window to receive a liquid reward. **d**, Trials concluded with a target sound preceded by a variable number of distractors. **e**, Lever release within a target’s response window was classified as a hit. Lever releases during delays or go-cue periods preceding a target were classified as false alarms (FA) and led to the trial abortion and initiation of a new trial. Misses counted when monkeys failed to release the lever during the target’s response window.

The monkeys discriminated every auditory category proficiently (Supplementary Fig. 1a), achieving high overall performance (monkey V: 96% ± 2.96%; monkey X: 89.8% ± 5.61%; mean ± SEM). They also showed faster reaction times to hits than to false alarms (Wilcoxon signed-rank test, p < 0.001) (Supplementary Fig. 1b), suggesting uncertainty in releasing the lever after distractors.

**Supplementary Fig. 1.**
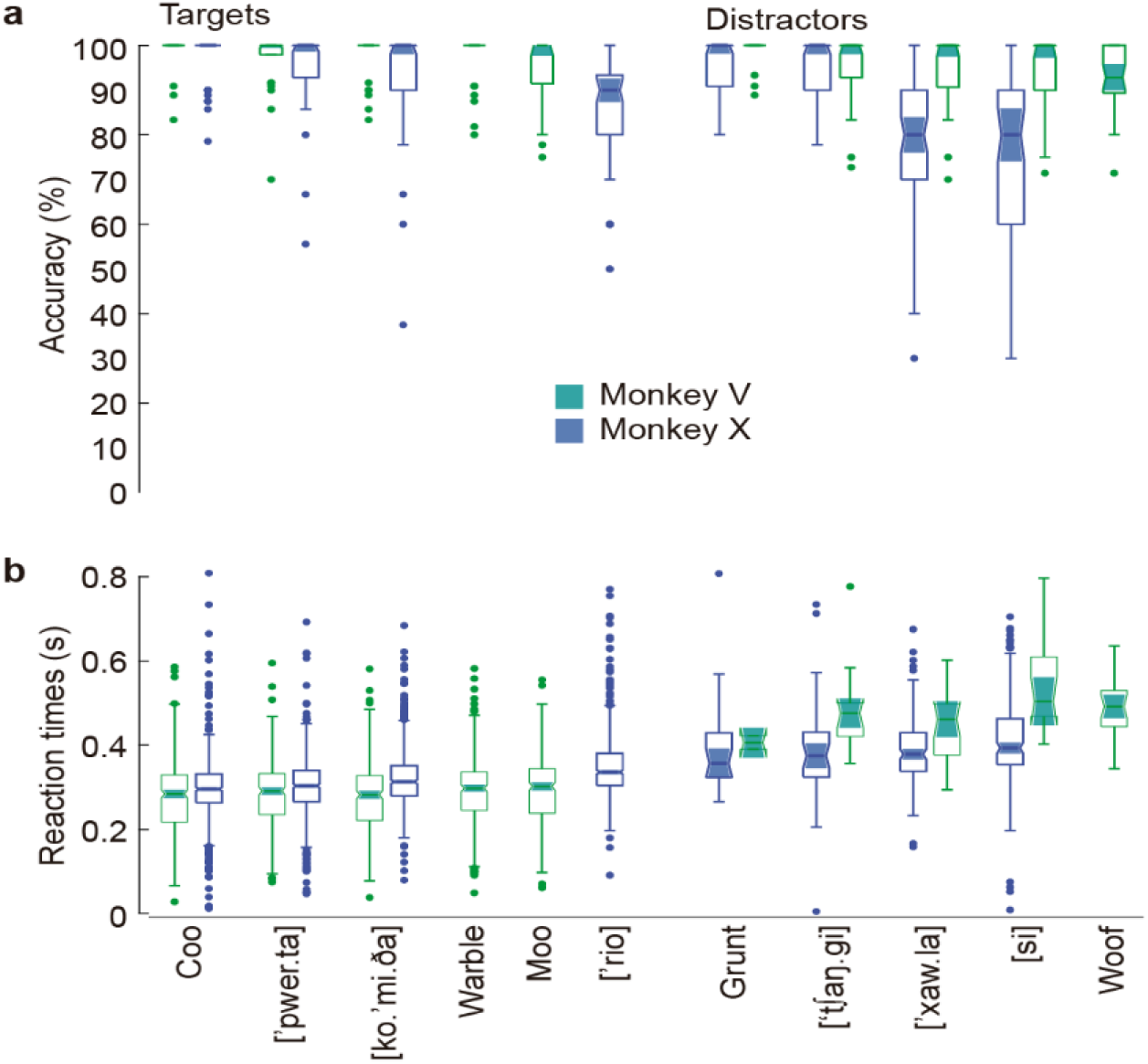
Behavioral performance. **a**, Monkeys performance in discriminating each sound (see Methods). Data points represent mean performance for individual sessions (Monkey V: n = 34 sessions; Monkey X: n = 63 sessions). The central bar indicates the grand mean, and error bars denote ± 1 SEM across sessions. **b**, Reaction times for lever releases after the visual go-cue following targets (hits) and distractors (false alarms). Principal Component Analysis (PCA) revealed robust engagement of the FPC neuronal population to the task (Fig. 2a, Supplementary Fig. 2a). PC1 showed pronounced modulations after the onset and offset of auditory periods, particularly for targets. PC2 captured the initiation of trials at the lever press and the reward periods in monkey V and tracked the first and last sounds in both monkeys. PC3 was associated with distractors in monkey V and with sound onsets in monkey X. Figure 2b (Supplementary Fig. 2b, c) shows that a significant proportion of FPC neurons responded to auditory stimuli (monkey V: 40.6%, 52/128; monkey X: 28.4%, 85/299; ZETA test, p < 0.05). Response latencies typically occurred within 200 ms of sound onset (Fig. 2c). In monkey V, the fastest mean latency was 111 ms ± 84 ms (mean ± STD) for [ˈpwer. ta], and the slowest was 174 ms ± 139 ms for a monkey warble. In monkey X, the fastest was 112 ms ± 105 ms for the pseudo-word [’ t∫aŋ. gi], and the slowest was 166 ms ± 120 ms for the word [si]. Some neurons exhibited early graded responses during auditory periods for targets and late graded responses for distractors, regardless of their position in a trial (Fig. 2d). Other neurons responded selectively to a few targets, distractors, or both (Fig. 2e).

Stimulus specificity of neural responses during auditory periods was assessed by projecting the neural response variability for each stimulus onto the PCA space. The results revealed a graded segregation of auditory categories along the first two principal components, which together explained approximately 40% of the variance in both monkeys (Fig. 3a). PC1 also distinguished targets from distractors during the auditory and delay periods in both subjects, indicating that the FPC encodes diverse auditory identities and their behavioral relevance.

A classifier trained to decode auditory categories out of the population activity showed that decoding accuracy exceeded chance at most auditory periods in monkey V, with two prominent peaks around 180 and 490 ms of auditory onset (Fig. 3b). Significant decoding in monkey X reached maximum accuracy around 220 ms. These results confirm that the FPC encodes diverse auditory categories. However, to determine whether decoding favored sounds within target or distractor groups, the classifier was trained on the sounds of each group separately. Decoding accuracy was observed in both groups (Fig. 3c). In monkey V, target identities were best decoded earlier (0.56 at 190 ms) than distractors (0.75 at 470 ms), whereas in monkey X, late decoding was observed in both groups (targets: 0.52 at 400 ms; distractors: 0.6 at 300 ms). This finding confirms that the FPC represents diverse auditory categories regardless of their behavioral association. To further assess whether the FPC also generated decision signals, the classifier was trained to distinguish between target and distractor groups regardless of the auditory categories they comprised. Figure 3d shows a peak accuracy achieved significantly after 150 ms of the visual cue onset in monkey V, in contrast to monkey X, where peak accuracy was reached during the delay period at 1260 ms after auditory onset. Collectively, these results establish that the FPC encodes both the identity of auditory stimuli during the auditory period and their associated behavior during the delay and visual periods.

We also conducted a clustering analysis to characterize how individual neurons contributed to encoding across different auditory categories. The results indicate that 80% of responsive neurons in monkey V and 79% in monkey X encoded different sounds during the auditory, delay, or visual task periods (Fig. 3e). While most neurons were selective for only one sound within a specific task period (Fig. 3f), many encoded two or more sounds, collectively representing all auditory categories (Fig. 3g). We then examined whether the neuronal encoding of multiple sounds was selective for targets or distractors, thus potentially reflecting decision-making activity. Here, most neurons encoded sounds from either group, but rarely synchronically. For instance, in monkey V, only 2% of bins in auditory epochs showed overlapping encoding, with similarly sparse overlap in other periods (delay: 0%; visual: 10%), and monkey X exhibited no instances of simultaneous encoding. These findings indicate that FPC single units primarily represent auditory categories more than encoding decision-making signals.

### Abstract coding of auditory categories

To determine if FPC representations are based on physical sound properties, we challenged the monkeys to discriminate morphs between a target and a distractor (e.g., between a monkey ’coo’ and a ’grunt’) mixed in different proportions (Fig. 4a). A subset of neurons responsive to the original learned sounds (9/20 in monkey V; 12/36 in monkey X) were also activated by the morphs. Figure 4b shows an example of a neuron responsive to several targets and distractors. Its latencies increased as the morph approached the original target sound, while its firing rate increased the farther the morph was from the learned target and distractor (Fig. 4c). The firing rates did not correlate with the morphs’ proportions despite this trend in the monkey’s behavior (Figs. 4d and e, respectively). Instead, the highest firing rates occurred at the highest coo-grunt mixture ratios, suggesting a role in detecting conflicting stimuli. At the population level, a classifier trained on a set of morphs significantly decoded them during the auditory period (Fig. 4f). However, only the 70% to 90% morphs were decoded for more than 5% of auditory period bins around the highest accuracy peak (Fig. 4g). These results suggest that FPC neurons create abstract representations of sounds rather than concrete representations of their physical properties.

To test whether these abstract representations remained perceptually constant to variations, we challenged monkey X with novel versions of the learned sounds (Fig. 5a). The monkey’s overall performance was 72.17% in the novel versions (Fig. 5b), compared to 90.47% in the corresponding learned sounds. Although behavior was partially affected by the detection of novel sounds, the animal successfully generalized. We recorded 33 neurons during the presentation of both learned and novel sounds. Of these, 16 were responsive at different task periods; 13 responded to learned sounds and 12 to the versions. However, neurons encoding novel sounds rarely encoded the corresponding learned categories (Fig. 5e).

A cross-decoding analysis was conducted to assess auditory constancy in the neuronal population. This involved decoding the novel versions using a classifier trained on the learned sounds and decoding the learned categories with the classifier trained on the novel versions. The results indicate that the population encoded either the learned or novel sounds, but not both (Fig. 5e), suggesting that although a few neurons exhibit constant sound encoding, the population represents learned and novel categories separately. Two examples illustrate this trend. First, a neuron that responded to learned targets and distractors increased its firing rate for novel sounds (Fig. 5f), indicating it could signal novelty.

Although that neuron maintained the representations of some categories (e.g., the monkey grunt), it also encoded novel sounds not previously represented in its learned representations, such as the coo vocalization and the [si] Spanish word. Second, an example of a neuron shifting its encoding of the learned [Ko.mi.da] and [Xau.la] sounds, to only encoding a novel version of the monkey grunt (Fig. 5g).

## Discussion

Auditory information is the primary sensory input to the frontopolar cortex (Burman et al., 2011; Petrides & Pandya, 2007; Barbas & Mesulam, 1985; Barbas et al., 1999; Hackett et al., 1999; Romanski et al., 1999a,b; Medalla & Barbas, 2014). Here, we examined the FPC involvement in auditory perception at the single-neuron level in two macaques trained to discriminate target sounds from distractors, including complex stimuli such as conspecific vocalizations and words (Melchor et al., 2021; Morán et al., 2021). Many FPC neurons responded robustly to one or more auditory categories and collectively represented the range of auditory stimuli (Fig. 2). Unlike previous studies of the neuronal activity during delay and feedback periods in visually guided tasks (Tsujimoto et al., 2010; Ferrucci et al., 2022; Nougaret et al., 2024), our findings show behaviorally relevant activity during auditory presentations, delays and visual-cue periods (Supplementary Fig. 2a). These results suggest a key FPC role in auditory perception, supporting the proposal that the FPC constitutes a frontal auditory field (Barbas & Mesulam, 1985; Medalla & Barbas, 2014).

**Fig. 2.**
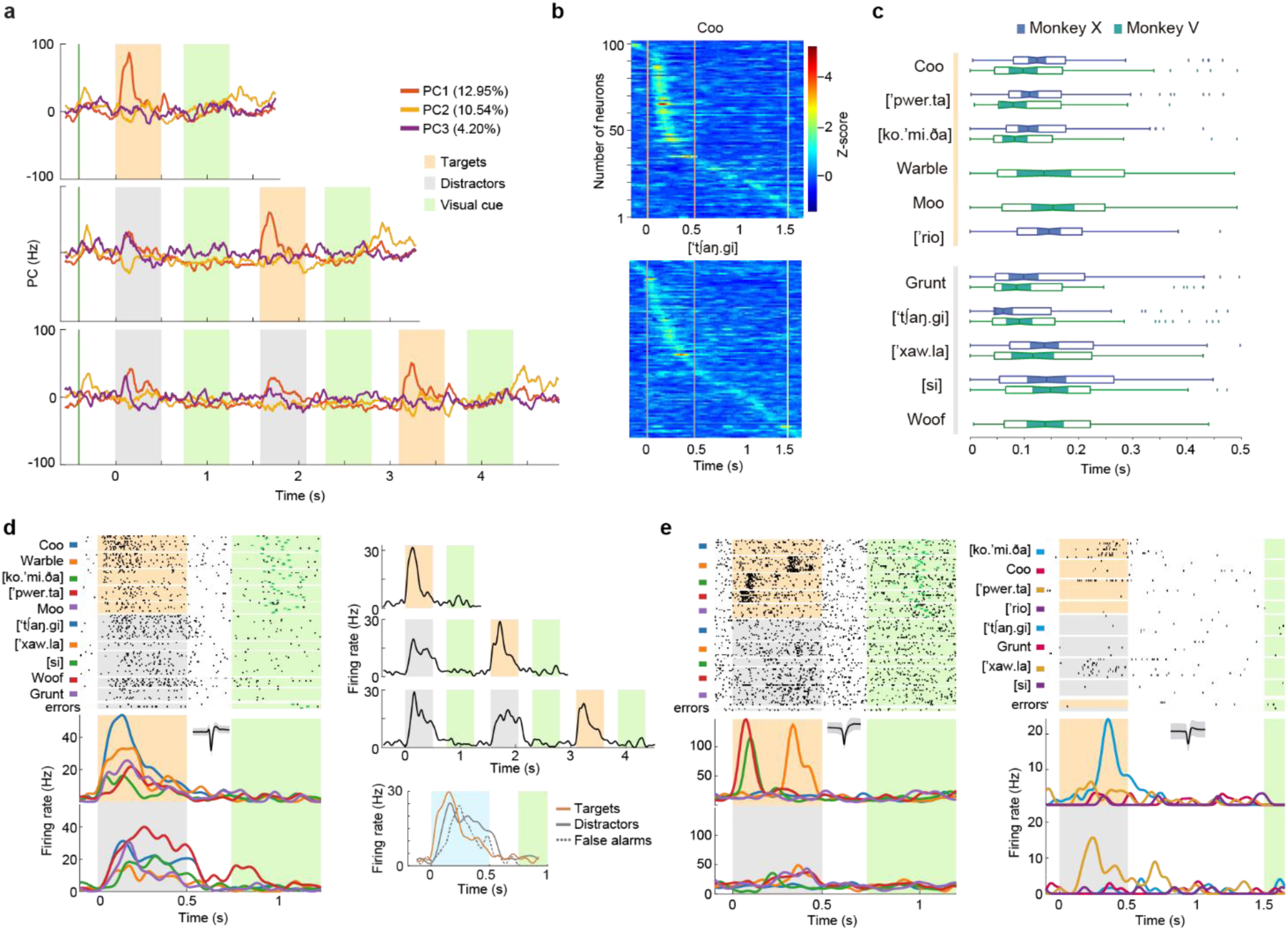
FPC neuronal responses during the auditory discrimination task. **a**, Temporal dynamics of the first three principal components of the neuronal population activity in monkey V. Activity is aligned to the onset of the first sound for trials with 1, 2, or 3 sounds. Green vertical line, lever press; orange, target sound periods; gray, distractor sound periods; green, go-cue periods. **b**, Heatmaps of mean firing rates in response to a target (coo) and a distractor (the Spanish word for ‘door’) for monkey X. Responses are shown for the auditory and delay periods for each sound, regardless of its serial position within a trial. Each row represents a recorded neuron, sorted by peak response time. The color scale indicates normalized firing rate. Orange or gray lines denote the onset and offset of sound; the green line marks the response window. **c**, Box plots of the neurons’ response latencies during the auditory periods for each target (orange group) and each distractor (gray group). **d**, Spiking activity of a neuron showing graded responses to targets and distractors. Upper left: Raster plot aligned to sound onset. Each tick is an action potential, and each row is a trial. Trials are grouped by auditory category (labeled at left). Orange, gray, and green boxes mark the targets, distractors, and visual go-cue periods. Lower left: Spike-density functions for each auditory category, color-coded as in the raster above. Inset: mean waveform modulation of the neuron ± 1 SD. Note the longer sustained responses to distractors compared to targets. Upper right: Peristimulus time histograms of the averaged activity of all targets and distractors in trials with 1–3 sounds. Same color code as in **a**. Lower right: mean firing rate comparison between targets, distractors, and false alarms during the auditory and delay periods, regardless of their position in a trial. **e**, Raster plot and spike-density function of a neuron reflecting abstract encoding of three targets labeled in colors as in **d**. **f**, Same format as **e**, but for a neuron showing abstract encoding of one target and one distractor.

**Supplementary Fig. 2.**
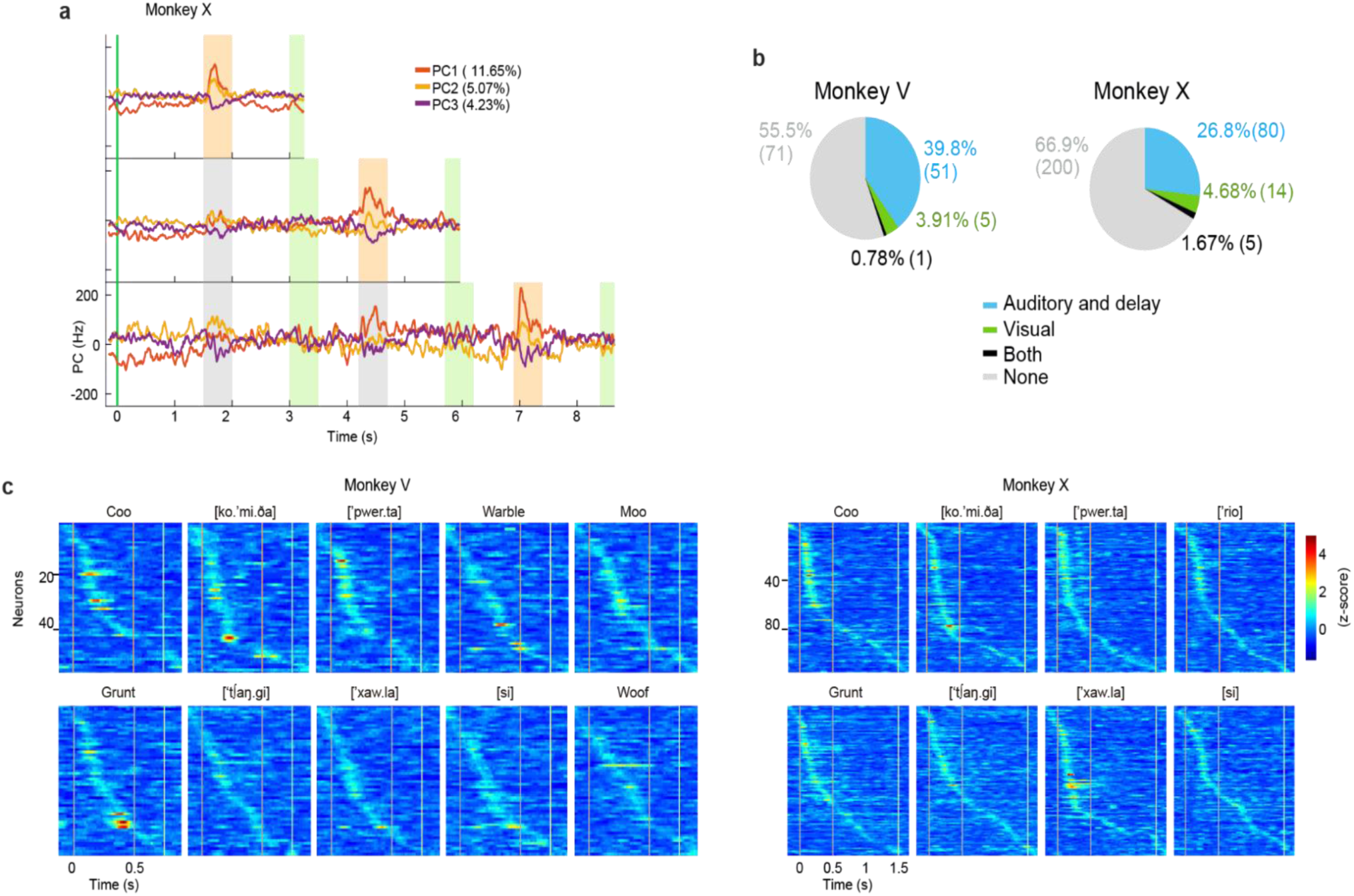
FPC neural population responses. **a**, PCA dynamics of FPC of population activity in Monkey X. Orange boxes mark target presentations, gray boxes, distractors, and green the visual go-cue. The green vertical line indicates the lever press. **b**, Proportion of FPC neurons (Monkey V, n = 128; Monkey X, n = 299) exhibiting significant stimulus-evoked responses during the auditory, delay, and go-cue periods (ZETA test, p < 0.05). **c**, Heatmaps displaying the firing rates of all recorded neurons in response to each target (top) and distractor (bottom) for Monkey V (left, n = 57 neurons) and Monkey X (right, n = 99 neurons). Activity is aligned to the auditory periods, regardless of their serial position within a trial. For each monkey, neurons are ordered (top to bottom) by the latency of their peak response. Firing rates are Z-scored relative to a pre-stimulus baseline period. Vertical orange or gray lines indicate the onset and offset of sound presentations, and the green vertical line indicates the go-cue onset.

Within the auditory processing hierarchy, the superior temporal gyrus (STG) and ventrolateral prefrontal cortex (VLPFC) fulfill distinct and well-established roles. STG neurons in the belt and parabelt regions respond selectively and with short latencies to vocalizations, thereby encoding their perceptual identity (Russ et al., 2008; Tian et al., 2001; Rauschecker et al., 1995; Tian & Rauschecker, 2004). STG can extract acoustic particularities from sounds (Russ et al., 2008). In contrast, the VLPFC operates at a more abstract level (Gifford III et al., 2005) but with a diminished capacity to distinguish subtle acoustic differences (Cohen et al., 2006). Neurons encoding vocalizations and words have been identified in both regions (Russ et al., 2008; Tian et al., 2001; Tsunada et al., 2011; Leonard et al., 2024; Romanski & Goldman-Rakic, 2002; Romanski et al., 2005; Plakke et al., 2013; Cohen et al., 2007; 2009; Russ et al., 2008).

The FPC processes learned vocalizations and words at a higher level of abstraction, using mechanisms distinct from those of the STG and VLPFC. Reports of neuronal latencies (Kajikawa et al., 2015; Nishimura et al., 2018; Kikuchi et al., 2014; Camalier et al., 2012) suggest that FPC response latencies (Fig. 2c) reflect direct anatomical inputs from auditory regions (Burman et al., 2011; Petrides & Pandya, 1988; 2007; Barbas & Mesulam, 1985; Barbas et al., 1999; Hackett et al., 1999; Romanski et al., 1999a,b; Medalla & Barbas, 2014), likely transmitted via the uncinate fasciculus and extreme capsule (Petrides & Pandya, 2007). These latencies are similar to those observed in the VLPFC (Romanski & Hwang, 2012), suggesting parallel processing. Functionally, most single FPC neurons are selective for a single auditory category (Fig. 3f), similar to the VLPFC (Romanski et al., 2005). The FPC population can decode auditory identities (Fig. 3e) as efficiently as the STG and more effectively than the VLPFC (Russ et al., 2008). Notably, FPC neurons collectively represent the entire set of sounds (Fig. 3g), consistent with evidence that item sets in humans drive FPC activation (Sakai, 2008; Pollmann, 2016).

**Fig. 3.**
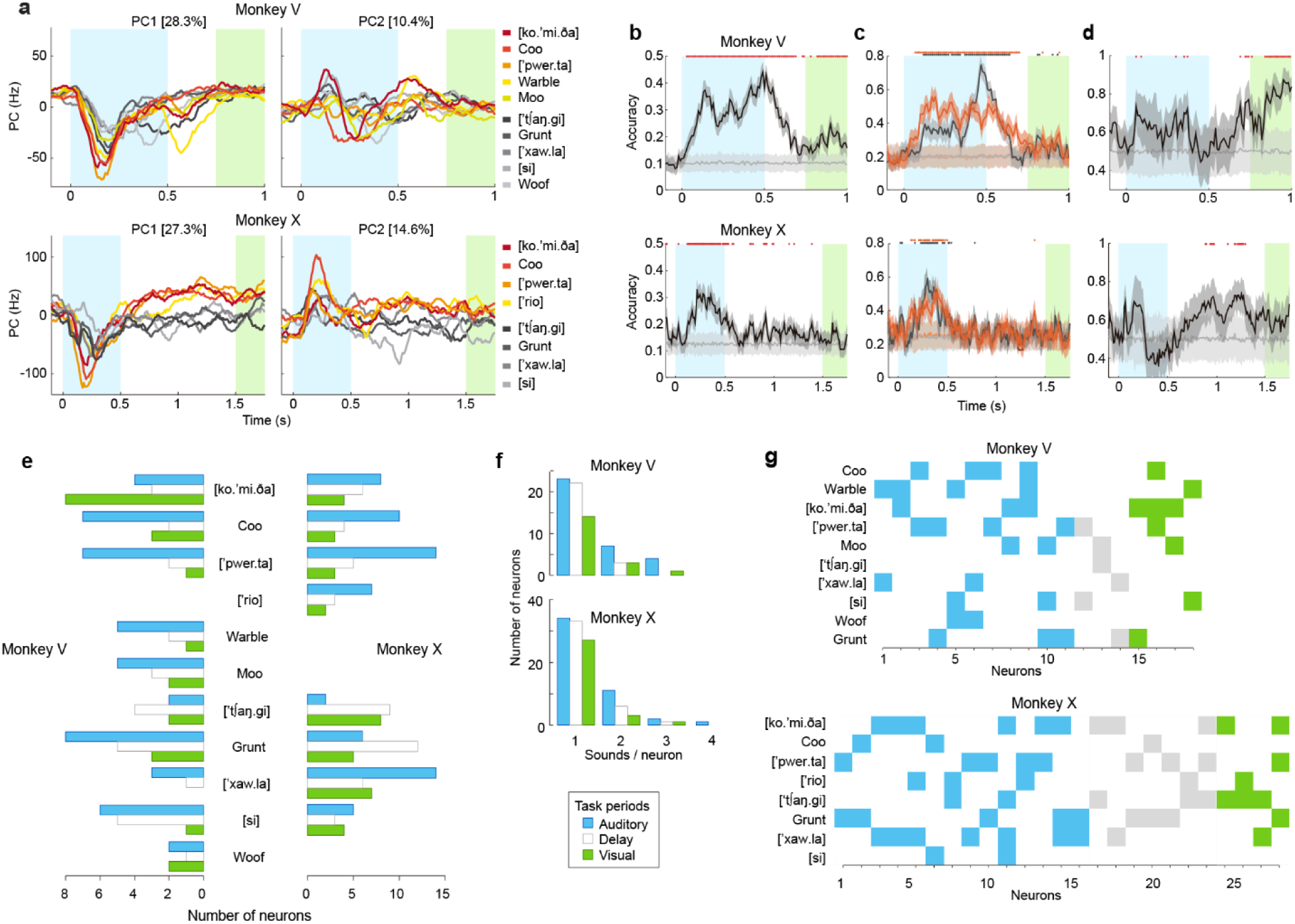
FPC population encoding of auditory categories. **a**, Dynamics of the first (left) and second (right) principal components of the neuronal population. Percentages indicate the variance explained by each component. Red hues denote targets, gray shadings, distractors. PC1 shows early tuning to auditory categories during the auditory epoch for both monkeys and later differentiates targets from distractors in monkey X, especially during the delay period. **b**, Time-resolved classification accuracy for individual sounds (see Methods). The shadings correspond to ±1 SD in cross-validation iterations. Horizontal lines indicate chance level (10% for monkey V, 12.5% for monkey X). The cyan and green boxes correspond to the auditory and go-cue epochs, respectively. The above color points indicate windows with accuracies above chance (p < 0.05). **c**, Decoding accuracy for individual sounds within the target (orange) and distractors (gray) groups. **d**, Decoding accuracy for discriminating between target and distractor groups. **e**, Count of neurons significantly coding for the different auditory categories in each monkey (p < 0.05, cluster-based permutation test). Colors correspond to the task events as in **a**. **f**, Number of single neurons encoding one or more sounds during distinct task epochs (p < 0.05, cluster-based analysis). Colors are the same as in **a**. **g**, Single neurons that encoded two or more sounds during the auditory (blue), delay (gray), or visual go-cue (green) periods.

The FPC’s abstract, non-parametric responses to auditory categories may function as a central hub for integrating auditory categories with task rules and goals. When we exposed the monkeys to auditory morphs created by gradually mixing a target and a distractor, responses did not vary with the acoustic proportions (Fig. 4), unlike those of SMA neurons (Morán et al., 2021). Instead, responses were selectively enhanced for the most ambiguous morphs, particularly those near 50% target-distractor similarity. Neuronal firing was highest for 50% morphs, lower for morphs with fewer target-like features, and lowest for morphs with higher target proportions (Fig. 4c, e). This pattern suggests that FPC neurons may contribute to explore-exploit behaviors (Mansouri et al., 2014; 2017; Zajkowski et al., 2017; Boorman et al., 2009; Gilbert et al., 2006; Koechlin et al., 1999) by assessing cost-benefit trade-offs (Tsujimoto et al., 2011; Mansouri et al., 2014; 2017; Koechlin et al., 1999). In this paradigm, the cost of missing a reward by producing a false alarm to a morph, which always appeared first in a trial, was higher than the cost of waiting for a definitive target, which always appeared last. Waiting for the second sound increased the probability of reward, whereas an immediate release resulted in a longer timeout before the subsequent trial. These findings are consistent with studies in humans demonstrating that FPC activates during challenging decisions (Tsujimoto et al., 2011; Mansouri et al., 2014, 2017; Koechlin et al., 1999) and with monkey studies showing that FPC ablation improves performance on high-conflict tasks (Mansouri et al., 2014).

**Fig. 4.**
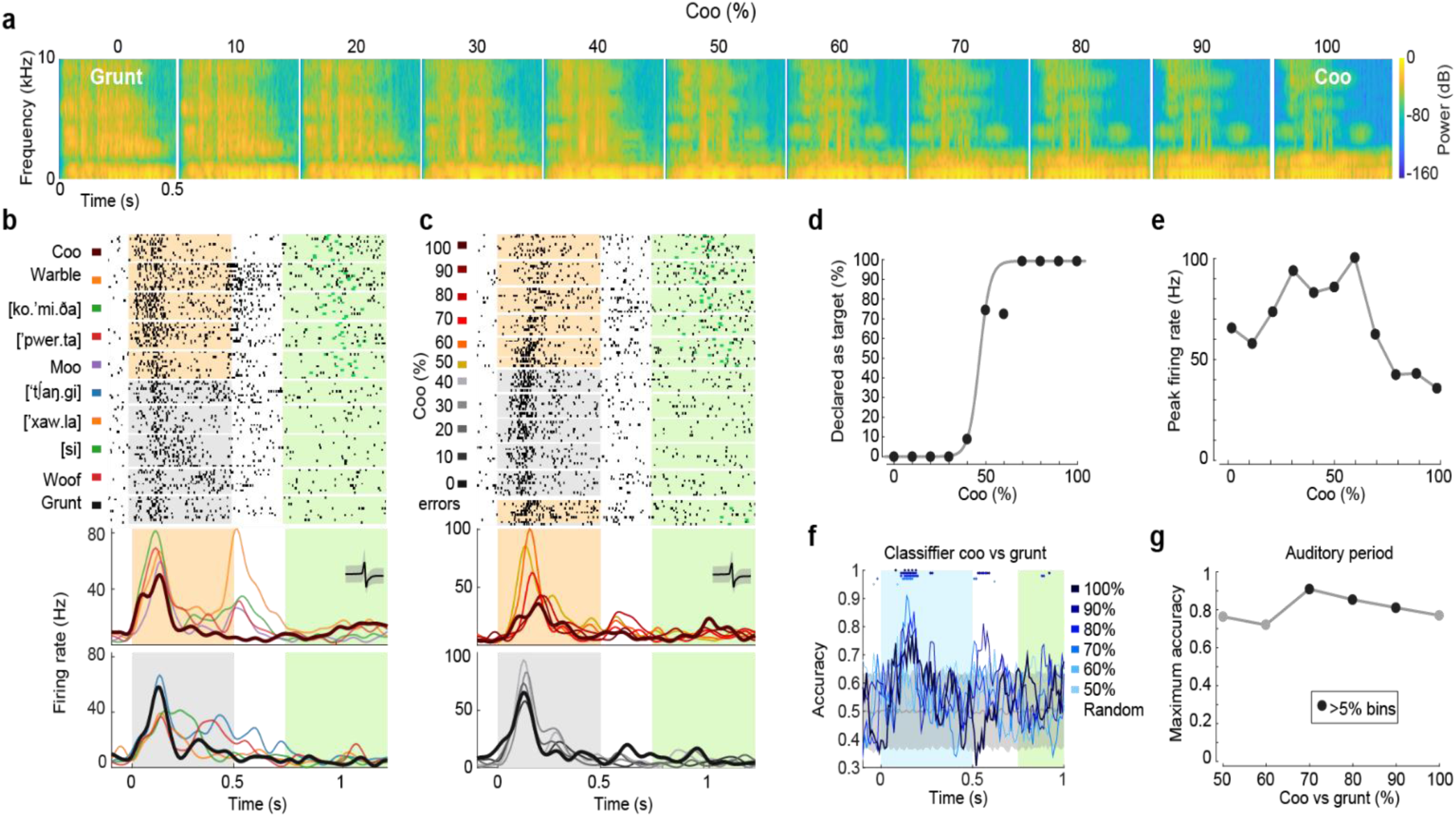
Abstract representation of sounds in the FPC. **a**, Target-distractor morph continuum. Spectrograms depict a 10% incremental acoustic transformation from a 100% grunt vocalization (distractor) to a 100% coo vocalization (target). **b**, Raster plot and spike-density functions of an example neuron’s responses to all learned target and distractors. Bold traces in the spike-density plots indicate the 100% coo (target) and 100% grunt (distractor). **c**, Responses of the neuron in b (see the matching neural waveforms at insets in lower panels) to the morph continuum. Bold traces in the lower panels represent 100% sounds highlighted in b. **d**, Discrimination of morphs by monkey V. Data points indicate the proportion of trials in which each morph was reported as a target. **e**, Peak firing rate of neuron in c, in response to each morph. Elevated firing is observed for intermediate morphs (∼50%), with the lowest response near the 100% target. **f**, Time-resolved classification accuracy in monkey V distinguishing neural population responses to different morphs. Shaded regions indicate time points at which decoding performance was significant (p < 0.05), specifically for morphs with target-similarity above 70%. **g**, Maximum classification accuracy observed in the analysis in f. Black circles indicate time bins where significant decoding was sustained for more than 5% of auditory bins.

When examining FPC responses to novel sounds that are acoustically similar to learned categories, some neurons increased firing to specific novel versions (Fig. 5f), while others switched to represent categories different from those initially encoded (Fig. 5g). Unlike VLPFC units responses to various sounds of same meaning or associated value (Gifford et al., 2005), or human hippocampal neurons showing invariant responses across stimuli of a single category (Quiroga et al., 2005), and neurons in the anterior temporal lobe that exhibit voice-identity invariance (Giamundo et al., 2025), FPC neurons do not display strong invariance. Instead, they respond differently to learned and novel stimuli (Fig. 5c), suggesting that the FPC supports flexible, context-dependent representations. One possible function of neurons that switch representations is the detection of analogies and semantic associations, as observed in human studies (Klein et al., 1995; Braver & Bongolatti, 2002; Green et al., 2006, 2010; Bunge et al., 2005, 2009). For instance, in the present task, animals were trained to associate auditory categories with specific actions, as in semantics, where sounds are linked to meanings. All target categories corresponded to “release” (the lever), while distractors corresponded to “hold,” with “lever release” serving as an analog for “reward.” The results show that the FPC associates sounds with hold-and-release actions (Fig. 3d) by delaying decision-making signals in the FPC population, perhaps for downstream computations. From an evolutionary perspective, the human FPC expansion (Ongur & Price, 2000; Semendeferi et al., 2001; Tsujimoto et al., 2011; Mansouri et al., 2017) may have enabled flexible object coding in FPC to support processing of acoustic variations that are essential for language and complex cognition.

**Fig. 5.**
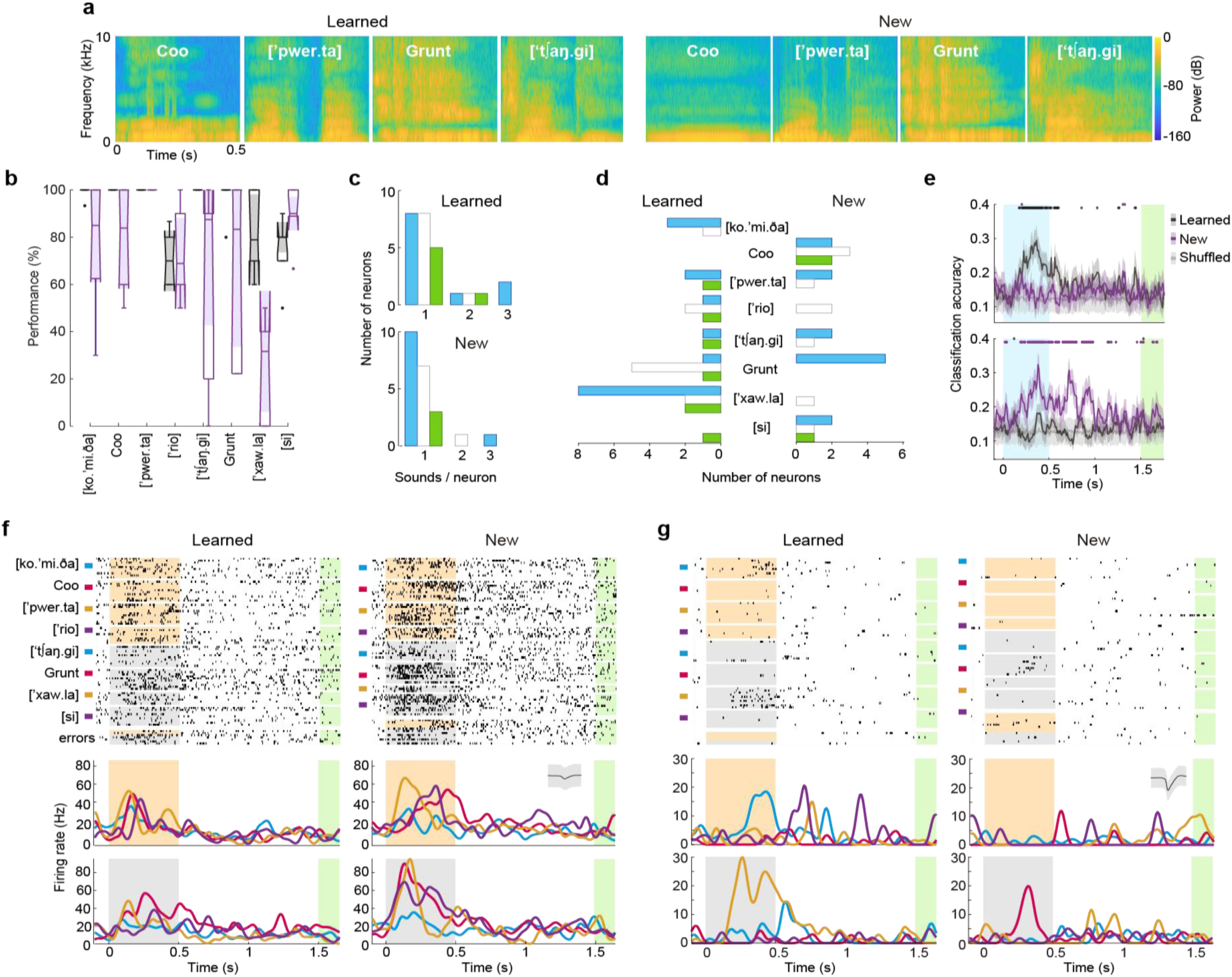
FPC detect novelty. **a**, Spectrograms of representative learned sounds and their corresponding novel versions, each produced by a different vocalizer, used to assess encoding invariance. **b,** Discrimination performance of monkey X for learned (black) and novel (purple) sounds. **c**, Number of neurons encoding one to three sounds (learned or novel) during distinct task epochs: auditory, delay, and go-cue. **d**, Number of neurons selectively encoding individual learned versus novel sounds. **e**, Cross-speaker neural decoding. Top: Accuracy of a decoder trained on responses to learned sounds and tested on novel versions (invariance test). Bottom: Decoding accuracy when the decoder was trained on novel versions and tested on learned sounds (novelty test). The gray line indicates the chance level. Shading represents ±1 SD across cross-validation iterations; colored dots above indicate time windows with significant decoding (p < 0.05). **f**, Neural responses to learned and novel sound versions. Left: Raster plot (top) and peristimulus time histogram (PSTH, bottom) for responses to original learned sounds. Right: Corresponding plots for novel versions. Inset: waveform of the recorded neuron. **g**, Raster (top) and PSTH (bottom) of a neuron responding to a novel sound category, which was different from the learned categories encoded by the same neuron.

Finally, the lack of prior reports on these findings in the FPC may be due to several factors. Single-unit recordings are not performed in the human FPC for clinical reasons, such as investigating epileptic foci or tumors. As a result, direct neuronal recordings are limited to non-human primates, the only species with FPC (Preuss & Goldman-Rakic, 1991). Still, neurophysiological recordings in these species are technically challenging (Mitz et al., 2009). Furthermore, most studies have used visual paradigms (Tsujimoto et al., 2010; Ferrucci et al., 2022; Nougaret et al., 2024), which do not match the FPC’s auditory specialization, but probably are a consequence of the belief that monkeys have difficulty with complex auditory tasks (Fritz et al., 2005; Ng et al., 2009), a belief that has only occasionally been challenged (Lemus et al., 2009a; 2009b; 2010; Morán et al., 2021; Melchor et al., 2021).

To summarize, this study provides the first direct evidence of neuronal mechanisms in the frontopolar cortex that maintain temporally organized auditory category maps and integrate relevant information to guide flexible actions. Future experiments would determine whether the neuroanatomical specificity of the FPC sunserves its broad functional repertoire, ranging from basic auditory discrimination to complex cognitive processes such as analogical thinking, cost-benefit analysis, explore-exploit behavior, and semantic association (Green et al., 2006; 2010; Bunge et al., 2005; 2009; Tsujimoto et al., 2011; Mansouri et al., 2014; 2017; Koechlin et al., 1999; Zajkowski et al., 2017; Boorman et al., 2009; Gilbert et al., 2006; Klein et al., 1995; Braver & Bongolatti, 2002;27, 28; Liu & Perfetti, 2003).

## Methods

### Ethics Statement

All procedures were in accordance with the Official Mexican Norm for the Care and Use of Laboratory Animals (NOM-062-ZOO-1999) and approved by the Internal Committee for the Care and Use of Laboratory Animals from the Institute of Cellular Physiology, UNAM (CICUAL-LLS200-22).

### Subjects

Two rhesus macaques (*Macaca mulatta*) participated in the experiments: Monkey V: 15-year-old male (10 kg), and Monkey X: 12-year-old female (6.6 kg). Both animals were housed individually in a room maintained on a 12-hour light/dark cycle, with filtered air and a constant temperature of 22 °C. The diet consisted of pellets, fruits, and vegetables, and a daily enrichment was provided through foraging toys, television (up to 3 hours per day of cartoons and wildlife videos), and access to a shared playground for climbing and socialization. Animals also had the opportunity to groom daily with conspecifics. Health status was monitored regularly by a veterinarian, and weight was tracked. To motivate task performance, water was restricted (10–12 hours) from Monday to Friday, ensuring a minimum daily intake of 20–30 mL/kg. Water was available *ad libitum* on weekends.

### Experimental Setup

The monkeys were head-fixed in a primate chair inside a soundproof booth. A metal lever (Crist Instruments Inc.) positioned at waist height was used to capture behavioral responses. Visual stimuli were displayed on a 21-inch color LCD monitor (1920 × 1080 resolution, 60 Hz refresh rate) placed directly in front of the animal. Auditory stimuli were presented at approximately 65 dB SPL via a Yamaha MSP5 speaker mounted above and behind the monitor (0.05–40 kHz), while a Logitech Z120 speaker positioned below provided background white noise at approximately 55 dB SPL.

### Behavioral Task

The animals were trained on an auditory discrimination task, as previously described (Melchor et al., 2021; Morán et al., 2021). In each trial, a monkey initiated the task by pressing and holding a lever in response to a gray circle displayed on the center of the screen. Following an initial delay of 0.5–1.5 seconds, a sequence of auditory stimuli was presented. For Monkey V, sequences consisted of 1 to 3 stimuli (each, 0.5 seconds in duration), while for Monkey X, sequences included 1 to 8 stimuli. In all cases, the target sound was presented only in the final position of the sequence. Each sound was followed by a delay (0.3–0.6 seconds for Monkey V; 1 second for Monkey X) and then a visual go-cue period. The go-cue was indicated by the central circle turning green for 0.5 seconds, initiating a 1-second response window. If the stimulus consisted of a target, the monkey was required to release the lever within this window to receive a juice reward. Late releases were classified as misses and were not rewarded. If the stimulus was a distractor, the monkey was required to continue holding the lever (a correct rejection). Releasing the lever during a distractor period was considered a false alarm, which led to the abortion of the trial and initiated a new trial without reward. The task was programmed using LabVIEW 2014.

### Auditory Stimuli

Auditory stimuli included monkey vocalizations and human words recorded in the laboratory. Additional animal vocalizations were obtained from the Freesound online library (https://freesound.org/). All sounds were sampled or resampled at 44.1 kHz, normalized to -10 dB RMS amplitude, and set to a duration of 0.5 seconds. Morph stimuli were generated as linear spectrotemporal interpolations between a target and a distractor sound using STRAIGHT software (Kawahara et al., 1999). For the sets of new auditory versions, each learned sound was substituted by a similar category emitted by different individuals. Each set comprised one novel sound per category repeated 10 times during the experiments.

### Surgical Procedure

Restrain headposts and Inox recording chambers (20 mm diameter for Monkey V, 10 mm for Monkey X) were surgically implanted. Chambers’ location above the left BA10 was determined using individual MRI scans and a stereotaxic atlas (Saleem & Logothetis, 2012). All surgical procedures were conducted under aseptic conditions and general anesthesia. Preoperative atropine (0.025 mg/kg) was administered before induction with ketamine/xylazine (6 mg/kg and 0.6 mg/kg, respectively) for Monkey V, and Zoletil50 (2 mg/kg) with xylazine (0.6 mg/kg) for Monkey X. Postoperative analgesia (ketorolac, 5 mg/kg) was provided.

### Electrophysiological Recordings

Extracellular recordings were obtained using a 5-channel microelectrode array (Thomas Recording) for independently movable electrodes (1–3 MΩ). Signals were acquired at 40 kHz using a 16-channel OmniPlex system (Plexon Inc.). The recording epochs differed between monkeys to align with the experimental objectives. For Monkey V, continuous activity was recorded from the pre-stimulus period through the behavioral response (lever release). For Monkey X, recordings were segmented around individual sounds, beginning 200 ms before sound onset and extending to the release of the lever or the end of the response window following correct rejections and misses. For direct comparisons, analyses were standardized to include activity within these periods up to the first 250 ms after the go-cue onset.

### Data Analysis

We used Offline Sorter from Plexon, and Kilosort 2 (Pachitariu et al., 2016; https://github.com/jamesjun/Kilosort2) to perform offline sorting of single neurons. Automated outputs were manually curated using Phy (https://github.com/cortex-lab/phy) and a custom interface. This curation included removing templates with noisy waveforms, splitting merged clusters, and discarding units that violated the refractory period. Artifacts were rejected using the PhyWaveformPlugin (https://github.com/jiumao2/PhyWaveformPlugin/tree/master). From the final curated pools (128 units from Monkey V and 299 from Monkey X), responsive neurons were identified using the ZETA test (https://github.com/JorritMontijn/ZETA, Montijn et al., 2021). Neural latencies were calculated within the ZETA framework using spike times from -200 to +500 ms relative to stimulus onset. Firing rates were obtained using two complementary temporal resolutions. For high-resolution visualization of peri-stimulus time histograms (PSTHs), 50 ms time windows were advanced in 10 ms steps and smoothed with a Poisson kernel (λ = 5). For population-level analyses, a 200 ms sliding window (10 ms steps) was employed to account for a limited number of trials per condition (typically 10). Dimensionality reduction for task events was performed using the PCA framework (Kobak et al., 2016; https://github.com/machenslab/dPCA) using MATLAB’s built-in PCA function (as described in Morán et al., 2021). For each trial, the inputs included the neuron identity, the presented stimulus, and firing rate in bins of 50 ms.

### Neural Decoding

A Bootstrap Aggregated Random Forest classifier trained on neuronal activity with default hyperparameters was implemented using MATLAB. Decoding accuracy was assessed per time-bin for each stimulus using 10 hit trials per neuron (5-fold cross-validation; 100 iterations). Statistical significance (p < 0.05) was determined from shuffled bin distributions. We implemented decoding classification adapted from Sarma et al. (2016) to distinguish decision-making signals from auditory representations. The classifier was trained to distinguish targets from distractors, excluding all trials of one target and one distractor, and tested with the excluded trials (permutation test, p < 0.05).

### Cluster-Based Neural Encoding

A clustering approach was implemented to identify stimulus-specific neural representations across time. For each temporal bin, k-means clustering was applied, with the optimal number of clusters determined by the Calinski-Harabasz criterion. To evaluate the probability of trials from a given stimulus in a cluster was significant, the following hypergeometric cumulative distribution function was computed:

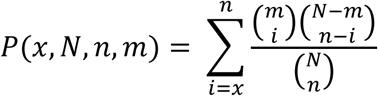

where N represents total trials in the set, n the trials of a specific stimulus, m the trials in a particular cluster, and x the observed co-occurrences between the stimulus trials and the cluster trials. Then, for each time-window, a Bonferroni correction was applied across cluster probabilities for each stimulus dividing the significance threshold (α = 0.05) by the number of clusters. A stimulus was considered significantly encoded if its probability fell below this corrected threshold.

To assess whether the clustering for a stimulus was significant throughout time windows, the entire clustering procedure was repeated 100 times to obtain a mean estimate of significant bins per stimulus, which was compared against the number of significant bins obtained from 100 clustering using randomized stimulus labeling (null distribution). Stimulus encoding was considered significant if the mean number of significant bins across iterations exceeded the 95th percentile of the null distribution.

## Acknowledgements

We thank Jonathan Melchor for assistance during experiments, Francisco Pérez from computing department of IFC, UNAM, and Manuel Ortínez and Aurey Galván from machine workshop from IFC (UNAM) for technical support. We also thank Professor Michael Brosch for valuable commentaries and suggestions.

## Authors’ contributions

Conceptualization: LL. Data Curation: MA, JV, TF. Formal Analysis: MA. Funding Acquisition: LL. Investigation: LL. Methodology: LL. Software: MA, TF, JV. Validation: JV, LL. Visualization: MA, LL. Writing original draft: MA, LL. Review & draft editing: JV, TF.

